# Microbiota-derived indole metabolites inhibit rotavirus infection *in vitro* and *in vivo* and in human infants

**DOI:** 10.1101/2025.10.29.685378

**Authors:** Nurul I. Wirusanti, Yannick van Schajik, Jongchan Kim, Josephine F. Frempong, Nagina Simkhada, Sydney Fisher, Nicholas Pucci, Goncalo J. Piedade, Charlie C. Luchen, Mwelwa Chibuye, Michelo Simuyandi, Caroline Chisenga, Suwilanji Silwamba, Kennedy Chibesa, Bertha T. Nzangwa, Dhvani H. Kuntawala, Sasirekha Ramani, Daniel Mende, Megan T. Baldridge, Bruno Sovran, Vanessa C. Harris

## Abstract

Rotavirus (RV) is the leading cause of life-threatening gastroenteritis in children under five despite effective RV vaccines availability, necessitating novel protective strategies for high-risk populations. Gut microbiota can modulate RV susceptibility and vaccine immunogenicity, yet underlying mechanisms are poorly characterized. Here, we demonstrate that microbiota-derived indole metabolites confer protection against RV infection. In healthy adults, higher fecal indole-3-acetic acid (IAA) and indole-3-propionic acid (IPA) levels associated with reduced fecal RV shedding following challenge. In human intestinal enteroids (HIEs), IAA pre-treatment inhibited human RV Wa G1P[8] replication via the aryl hydrocarbon receptor (AhR) pathway. In mice, treatment with AhR agonist indole-3-carbinol (I3C) significantly reduced fecal shedding of murine RV strain EDIM-Cambridge. Finally, Zambian infants with active RV infection exhibited lower fecal IAA and IPA levels than age-matched healthy controls. These findings demonstrate that microbiota-derived AhR ligands consistently inhibit RV infection across three experimental models and hold promise for protecting at-risk pediatric populations.

## Introduction

Rotavirus (RV) is the most important etiology of life-threatening gastroenteritis in children under-five, responsible for over one-third of diarrheal deaths globally^1^. There is a disproportionate burden of RV disease in low- and middle-income countries (LMICs), where over 90% of all RV-related deaths occur^2^. These disparities are likely driven by limited access to healthcare, a higher prevalence of comorbid conditions and concurrent enteric infections, alongside a reduced effectiveness of oral RV vaccines in countries with high RV mortality^3,4^. Given these challenges, novel RV prevention and treatment strategies are needed for children living in high-mortality settings.

Evidence from both human and animal studies to date have demonstrated that the gut microbiota can modulate RV infection and vaccine immunogenicity^5–22^. Currently licensed RV vaccines contain live attenuated viruses and must replicate in small intestinal epithelial cells to elicit protective immunity, mimicking natural infection^23^. Microbiome composition at the time of vaccination has been associated with fecal rotavirus vaccine shedding, a proxy for RV replication, in several observational studies in children from LMIC^11–13,15^. However, studies have had limited reproducibility, with bacterial taxa associated with RV shedding varying across geographic regions and populations^11–13,15^. In contrast, there has been limited assessment of whether the microbiota-derived metabolites and the functional capacity of the microbiota associate with RV infection or vaccine immunogenicity.

Microbiota-derived metabolites can alter host immunity by priming immune cell maturation and differentiation, modulating cytokine production, maintaining epithelial barrier integrity, and serving as an energy source for intestinal cells^24^. Further, numerous microbiota-derived metabolites, including short-chain fatty acids (SCFAs) and secondary bile acids, have been shown to regulate viral infections and disease outcomes^25,26^. These microbiota-derived metabolites can regulate the expression of viral receptors and co-receptors on host cells^27,28^, prime innate immune responses that are critical in virus infection, such as type I and III interferon (IFN)^29,30^, as well as control the induction of cytokines that convey signals to T or B cells ^31^, thereby determining host susceptibility to virus infection and modulating disease severity. In parallel with microbiota composition, microbial metabolites vary by geography and across human populations^32,33^ and may be a modifiable factor impacting host susceptibility to RV infection.

This study aims to investigate the role of microbiota-derived metabolites in modulating RV infection. Using samples from a previously published challenge study in healthy adults^7^, we found that microbiota-derived indole metabolites in the tryptophan pathway are associated with reduced RV fecal shedding. Dietary tryptophan can be metabolized either through host-dependent pathways, including the kynurenine and serotonin pathway, or through microbiota-dependent metabolism – the indole pathway^34,35^. Microbiota-derived indoles are well-known ligands for the aryl hydrocarbon receptor (AhR), a transcription factor that regulates mucosal immunity, barrier function, and epithelial homeostasis^34,35^. Indole metabolites and the AhR pathway have been implicated in protection against inflammatory bowel diseases^36,37^, metabolic diseases^38,39^, and enteric bacterial infections^40^. However, their role in enteric virus infections such as RV remains largely unexplored.

Using human intestinal enteroids (HIE) cultures and a murine model of RV infection model, we demonstrated that indole metabolites suppressed RV replication through an AhR-dependent mechanism. Finally, we validated these findings in a high-risk Zambian infant cohort, showing that RV-infected infants exhibit significantly reduced fecal levels of indole metabolites compared to non-infected infants. Together, our findings position microbiota-derived indoles and the AhR signaling axis as key modulators of RV infection and replication, with implications for microbiota-targeted interventions in the prevention and treatment of enteric viral infections.

## Results

### Fecal indole metabolites associate with reduced RV shedding in an adult challenge study

To identify microbial metabolites associating with RV replication, we leveraged fecal samples collected from a previously described randomized controlled trial in healthy adults, in which antibiotic-mediated modulation of the gut microbiota was associated with increased fecal RV shedding^7^. Briefly, a total of 63 healthy adult participants were randomized 1:1:1 ratio to receive either seven days of treatment with oral vancomycin (narrow-spectrum), a combination of oral vancomycin, ciprofloxacin and metronidazole (broad-spectrum) antibiotics or no antibiotics (control). Following a 48-hour washout period, all participants were challenged with a single dose of oral rotavirus vaccine (Rotarix^TM^, ORV) containing at least 10L 50% cell culture infectious dose (CCID₅₀) of the live attenuated human rotavirus G1P[8] strain. Fecal ORV shedding, as a proxy of RV replication, was monitored from day 1 to day 6 post-vaccination. Both narrow- and broad-spectrum antibiotic treatment increased the proportion of participants with detectable fecal RV shedding and elevated the concentration of fecal RV antigen, as compared to control^7^. Post-antibiotic treatment fecal samples were collected 48 hours following antibiotic cessation, just prior to ORV administration. In the present study, we performed metabolomics on the post-antibiotic fecal samples to identify fecal metabolites correlating with subsequent fecal RV shedding, measured in the 7 days following challenge **(Fig. 1a)**.

**Figure 1.**
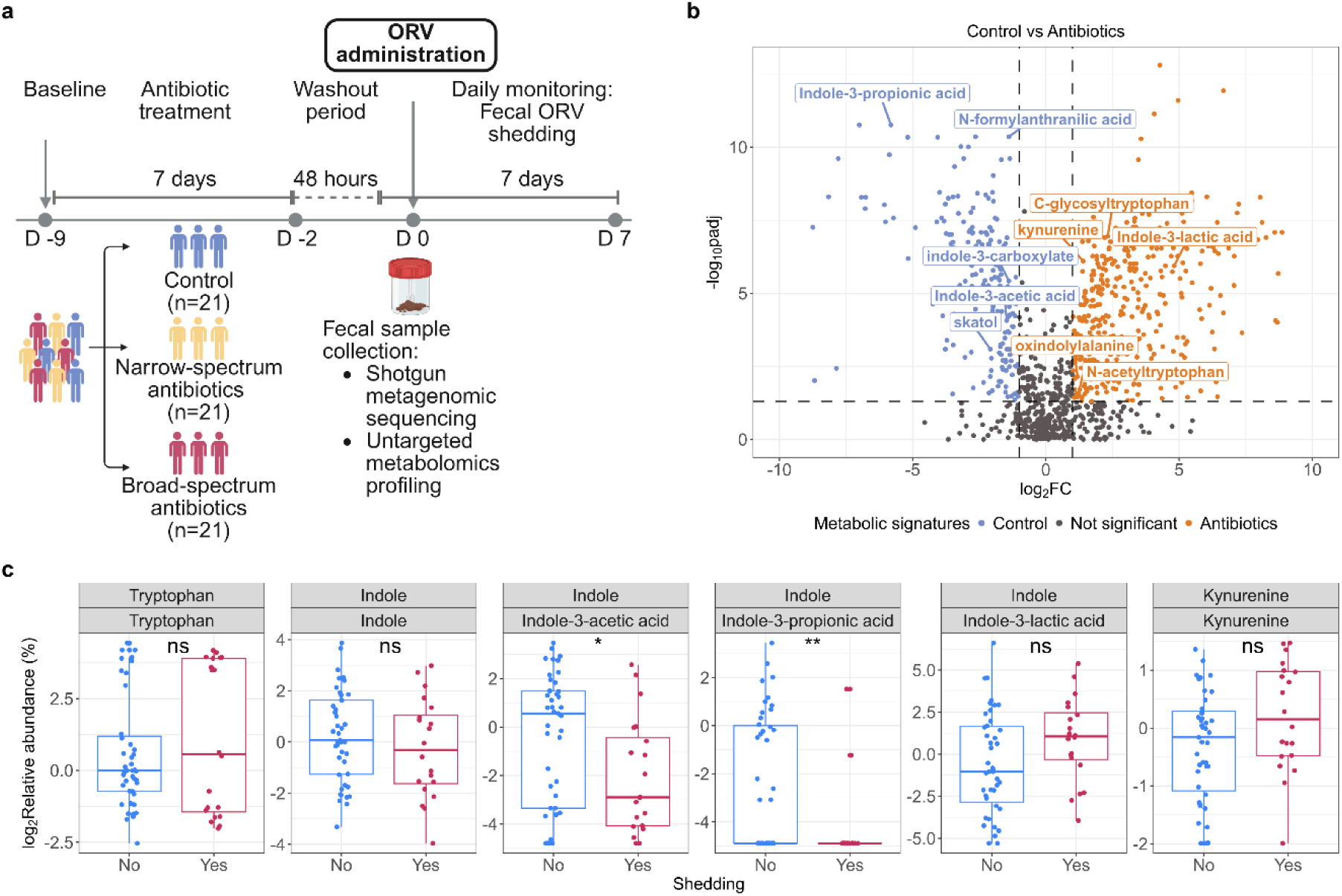
Fecal indole metabolites associate with reduced RV shedding in healthy adults challenge study. **(a)** Schematic overview of oral rotavirus vaccine (ORV) challenge study in healthy adults. **(b)** Volcano plot of differentially abundant fecal metabolites between control and all antibiotic-treated groups (log2foldchange >1 or <-1; p.adj <0.05, Mann-Whitney U test with FDR correction). Metabolites belonging to the tryptophan group are labeled. **(c)** Relative abundance of select tryptophan metabolites in RV shedders (n=20) and non-shedders (n=43). The bottom header of each panel shows the metabolite names, top header indicates the pathway classification (kynurenine or indole pathway). Boxplot indicates first, second and third quartiles of data, whiskers indicate maximum and minimum values. P-values are based on Wilcoxon test with FDR correction (ns: non-significant, * p< 0.05; ** p< 0.01; *** p< 0.001).

Differential abundance analysis showed that antibiotic treatment significantly altered tryptophan metabolism (log2foldchange >1 or <-1; p.adj <0.05, Mann-Whitney U test with FDR correction) **(Fig. 1B, Supplementary Fig. 1a-b)**. Microbiota-derived indole metabolites, including indole-3-acetic acid (IAA), indole-3-propionic acid (IPA), indole-3-carboxylate and skatole, were significantly enriched in the control group compared to the antibiotic-treated group **(Fig. 1b, Supplementary Fig. 1a-b).** Conversely, host-derived kynurenine was more abundant in antibiotic-treated subjects as compared to control **(Fig. 1B, Supplementary Fig. 1a-b**).

We next tested whether fecal metabolites associated with RV shedding status, irrespective of antibiotic treatment allocation. While fecal tryptophan levels did not differ between RV shedders and non-shedders, there was a significantly higher relative abundance of downstream indole metabolites, IAA and IPA, in subjects without RV shedding **(Fig. 1c).** Further, IAA and IPA were associated with lower concentrations and duration of RV shedding following ORV administration (area under curve; AUC) (**Supplementary Fig. 1d)**. Kynurenine tended to be higher in RV shedders, although the difference did not reach statistical significance **(Fig. 1c)**. All other metabolites in the indole and kynurenine pathways were not significantly different between shedders and non-shedders **(Supplementary Fig. 1c)**. Together, these data suggest that higher relative abundance of microbiota-derived indole metabolites at baseline are associated with reduced ORV shedding in healthy adults.

### Indole-3-acetic acid restricts RV replication in human intestinal enteroids

The observed negative association between the abundance of microbiota-derived indole metabolites (IAA and IPA) and fecal RV shedding in the adult challenge study suggested that indole metabolites may protect against RV infection by modulating viral replication. Since IAA is a potent AhR agonist^35^, we hypothesized that the action of IAA on RV replication was mediated through the AhR pathway. To test this hypothesis, we evaluated the impact of IAA on RV replication and compared it with host-derived kynurenine, using both adult and infant human intestinal enteroids (HIE). Host-derived kynurenine is also known as a potent AhR agonist^41^, and was included to examine distinctions between microbiota- and host-derived AhR ligands. Differentiated HIE monolayers were pretreated with 0.1% DMSO (vehicle), 50 µM kynurenine or 500 µM IAA for 24 hours, followed by RV infection (human G1P[8] WA strain) at multiplicity of infection (MOI) of 1 and incubation with the compounds for another 24 hours **(Fig. 2a)**. These concentrations, selected to reflect physiological concentration detected in the fecal samples, were confirmed to be non-cytotoxic to the HIE as measured by no significant changes in viability or LDH release compared to untreated controls **(Supplementary Fig. 2a-b)**. Consistent with the adult challenge study, IAA significantly reduced RV titer compared to vehicle **(Fig. 2b)**, with a more pronounced effect seen in the adult HIE **(Supplementary Fig. 2c)**. In infant HIE, neither kynurenine nor IAA reduced RV titers significantly relative to vehicle, however IAA-treated HIE showed lower RV titer compared to kynurenine-treated HIE (**Supplementary Fig. 2d)**. These findings indicate that IAA restricts RV replication in human intestinal epithelium.

**Figure 2.**
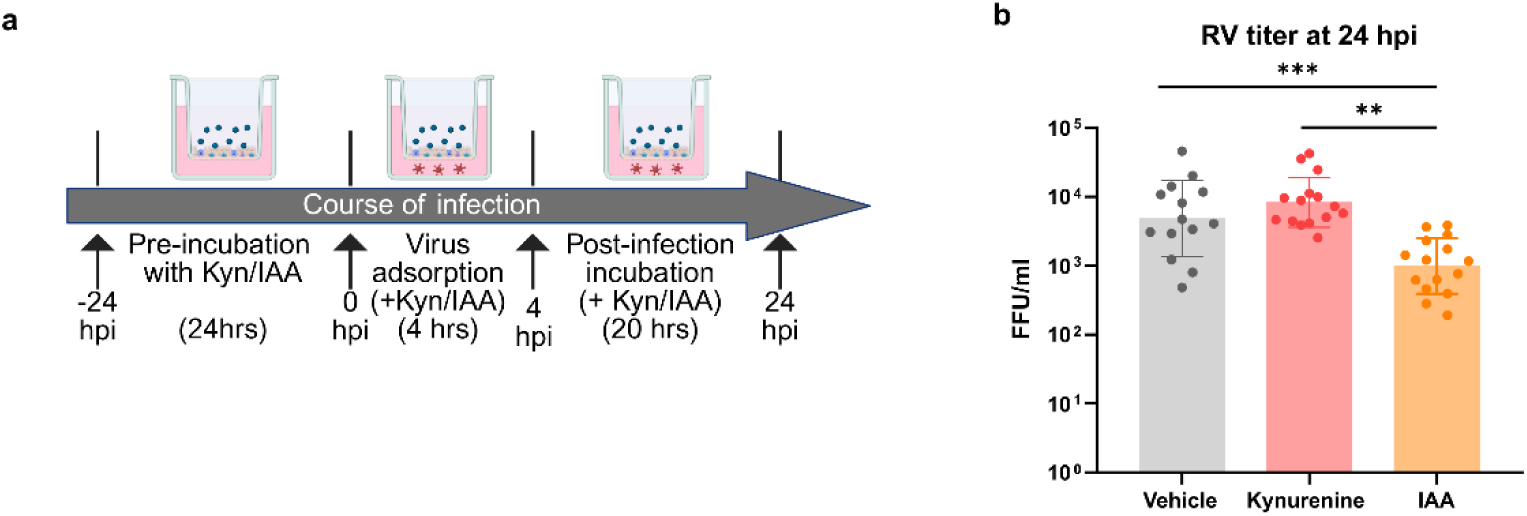
IAA inhibits RV replication in HIE. **(a)** Schematic overview of RV infectivity assay protocol in HIE. **(b)** Production of RV infectious virus particles with 0.1% DMSO (vehicle), 50 µM kynurenine or 500 µM IAA treatment, quantified at 24 hpi using the fluorescence focus assay (FFA). Data (geometric mean ± geometric SD) were collected from 3 adult and 3 pediatric HIE lines across 8 independent experiments. P-values are based on Kruskal-Wallis test with Dunn’s multiple comparisons test (* p<LJ0.05; ** p<LJ0.01; *** p<LJ0.001).

### Kynurenine and IAA activate AhR signaling in RV-infected HIE

To understand whether AhR mediates the effect of IAA on RV replication, and to characterize the distinct molecular pathways induced by IAA and kynurenine **(Fig. 2a, Supplementary Fig. 2a-b)**, we performed transcriptomic profiling via RNA-sequencing on adult and infants HIEs pre-treated with these compounds and subsequently infected with RV. Monolayers were collected 5 hours post-infection (hpi) from mock-infected and RV-infected HIEs for RNA extraction and gene expression analysis.

Principle coordinate analysis (PCoA) showed that the gene expression profiles differed primarily by age group (adult vs infant status) (Permanova p = 0.002), while treatment and infection status had modest effect (Permanova p = 0.337 and 0.557 respectively) **(Supplementary Fig. 3a-c).** Analysis of the top 10 differentially expressed genes (DEGs) revealed that RV infection alone significantly increased the expression of inflammatory markers including *Ifit1, Nfkb1a, Tnf, and Ccl20* **(Supplementary Fig. 3d)**, consistent with previous studies^42,43^. Notably, these genes were more strongly upregulated in adult HIEs than in infant HIEs **(Supplementary Fig. 3d)**. Similarly, IFN-stimulated genes (ISGs) were more highly upregulated in adult HIE compared to infant HIEs **(Supplementary Fig. 3e)**. Given these differences, we analyzed the infant and adult lines separately, with a focus on the infant-derived line given RV is a pediatric disease.

In RV-infected infant HIEs, pre-treatment with IAA induced AhR activation, marked by the upregulation of *Cyp1a1* expression **(Fig. 3a)** and enrichment of the cytochrome P450 pathway as compared to RV infection with vehicle **(Fig. 3b)**. IAA pre-treatment also induced the expression of *Oas1* **(Fig. 3a)**, an ISG downstream of the IFN pathway^44^ and enriched pathways known to influence RV replication, including prostaglandin receptor activity^45^ and bile acid binding^46^ **(Fig. 3b)**. RV infection with kynurenine treatment also induced AhR activation, evidenced by the upregulation of *Cyp1a1*, *Cyp1b1* and *Aldh1a3* expression **(Fig. 3c)**, along with enrichment of the P450 pathway **(Fig. 3d)**. Notably, RV infection with kynurenine treatment also induced the expression of MMP1 and enrichment for ferroptosis pathway as compared to control, which was not noted with IAA administration **(Fig. 3c-d)**. Importantly, RV infection alone did not markedly alter gene expression at 5 hpi **(Supplementary Fig. 3f-g)**, suggesting that at this early time point, observed transcriptional changes were primarily driven by AhR ligand treatment. These findings support AhR activation as a key mechanism through which microbiota-derived IAA suppresses RV replication in the gut epithelium.

**Figure 3.**
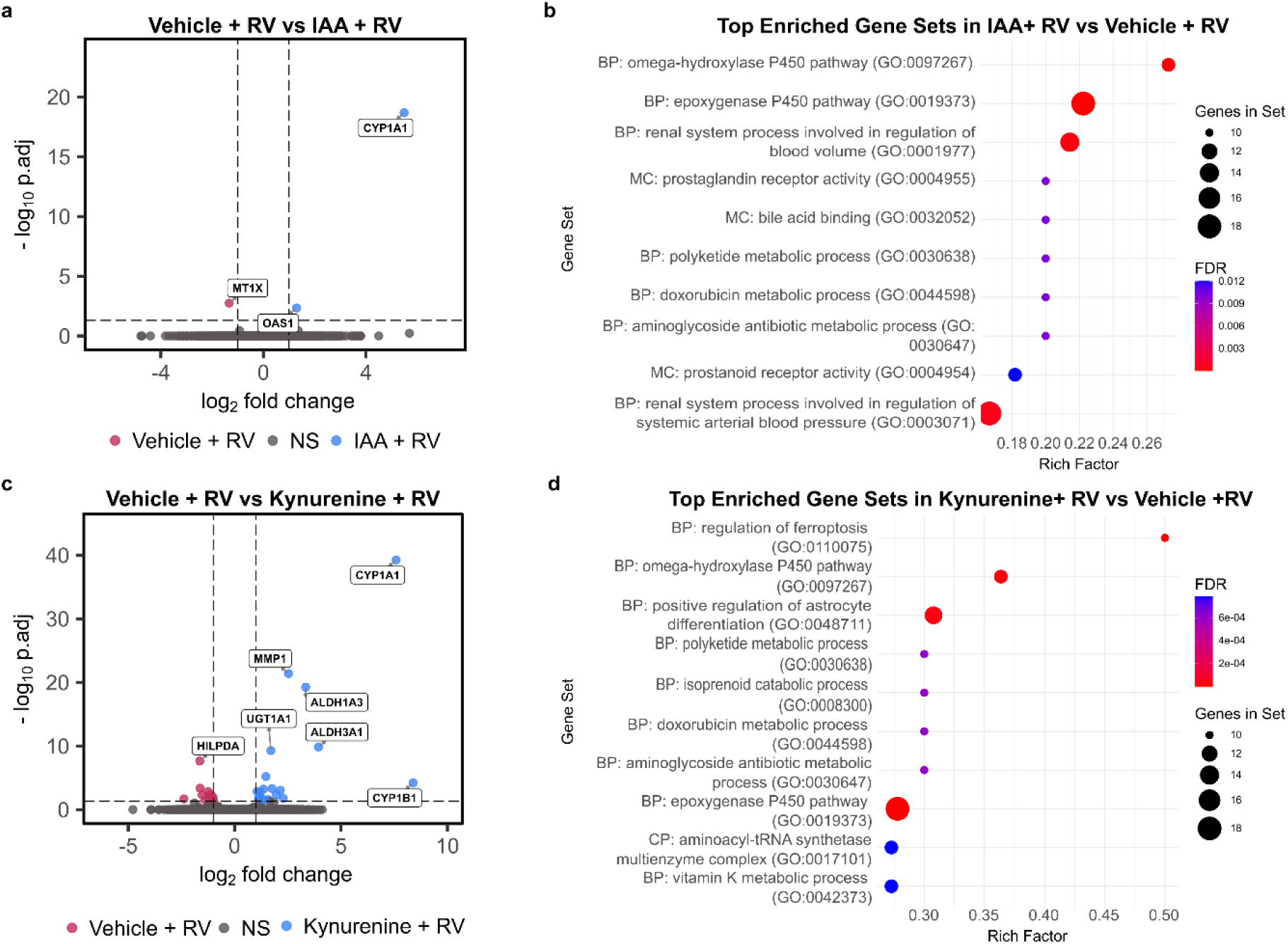
RNA sequencing analysis of RV-infected HIE pre-treated with IAA and kynurenine. **(a)** Volcano plot showing differentially expressed genes (DEG) compared between RV-infected IAA-treated HIE (n=2) and RV-infected vehicle-treated HIE (n=2), **(b)** Gene set enrichment analysis for IAA+RV compared to vehicle + RV based on comparison in (A), **(c)** Volcano plot showing DEGs compared between RV-infected kynurenine-treated HIE (n=2) and RV-infected vehicle-treated HIE (n=2), **(d)** Gene set enrichment analysis for kynurenine + RV compared to vehicle + RV based on comparison in (C). For (A) and (C) DEGs were defined as log2foldchange >1 or <-1; p.adj <0.05 (Wald test with Benjamini-Hochberg FDR correction). BP = Biological process, CC = Cellular component, MF = Molecular function.

### AhR agonist administration protects against RV infection *in vivo*

We next evaluated whether a known AhR agonist could similarly inhibit RV infection *in vivo*. C57BL/6 mice received daily oral administration of AhR agonist indole-3-carbinol (I3C), which has been previously administered in adult humans^47–49^. I3C administration began three days prior to murine RV (MRV) infection and continued to 12 days post infection (dpi). Fecal RV shedding was monitored daily from the 0-12 dpi. **(Fig. 4a)**. Fecal shedding began at 2 dpi and lasted till 5-6 dpi. I3C treatment led to an earlier resolution of RV shedding and significantly reduced viral loads during peak infection at 3-4 dpi **(Fig. 4b)**. Further, I3C treatment resulted in an overall lower level and duration of fecal RV shedding (area under curve; AUC) (p=0.0286) (**Fig. 4c)**. Intestinal tissues collected at 12 dpi revealed elevated *Cyp1a1*, *Ahr* and *Il22* gene expression in I3C+RV group, confirming AhR pathway activation **(Supplementary Fig. 4).** These results demonstrate that exogenous AhR activation suppresses RV replication *in vivo*.

**Figure 4.**
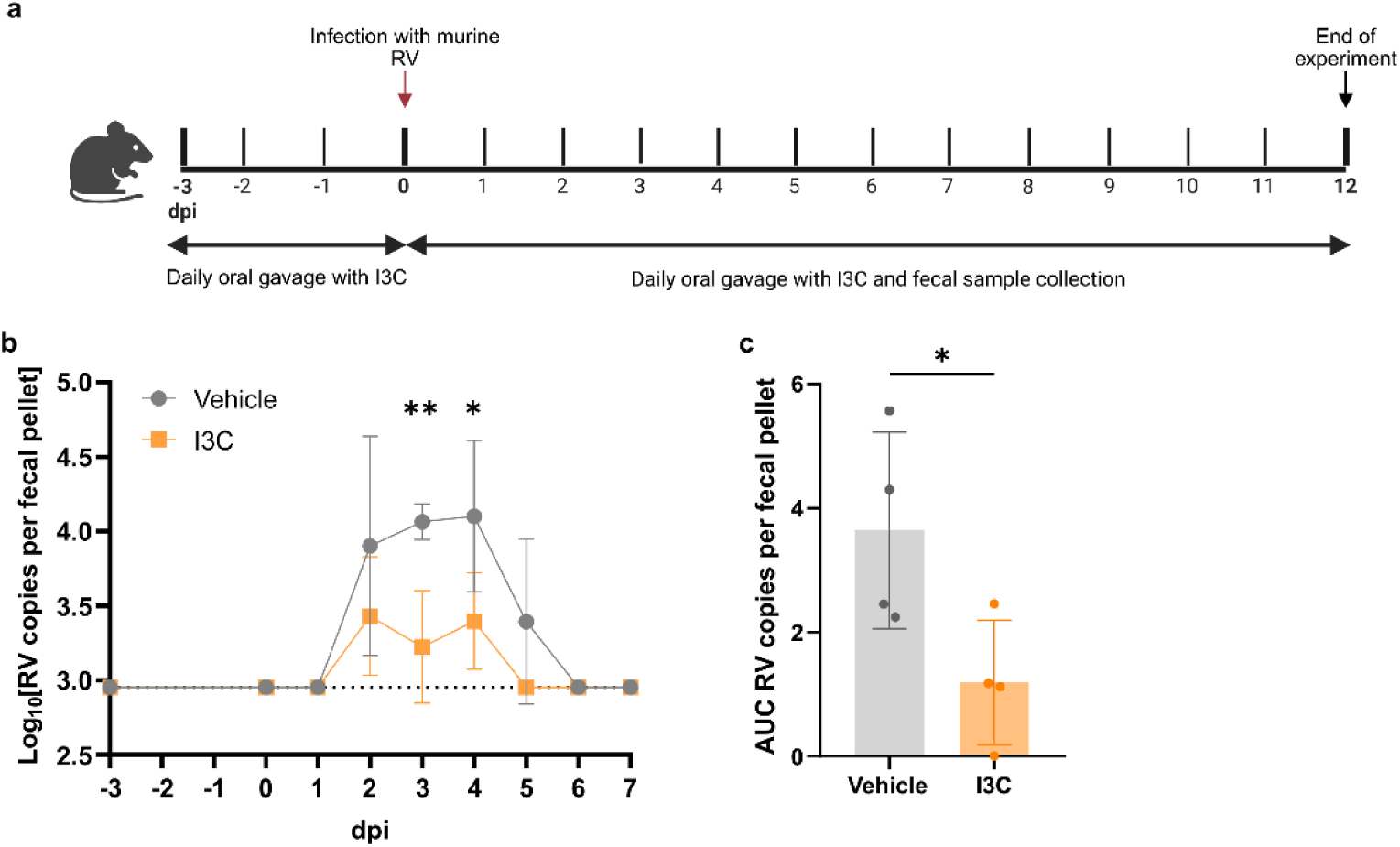
AhR agonist I3C reduces RV replication in C57BL/6 mice. **(a)** Experimental design of murine RV infections. **(b)** RV genome copies in fecal samples of RV-infected mice, with and without I3C treatment over time. **(c)** Area under curve (AUC) analysis of RV genome copies on 0-7 days post infection (dpi). Data (mean ± SD) are from 1 independent experiments with n=4 per group. P-values in (b) are based on 2-way ANOVA with Sidak’s multiple comparisons test. P-value in (c) is based on unpaired t-test (For both * p<LJ0.05; ** p<LJ0.01; *** p<LJ0.001).

### Zambian infants with RV infection have reduced fecal indole metabolites

Finally, to validate the clinical relevance of our findings in a relevant at-risk pediatric population, we analyzed fecal samples from infants with RV-confirmed gastroenteritis (n=35) enrolled in a diarrhea surveillance study in Zambia (NCT04312906 ClinicalTrials.gov) and compared their fecal tryptophan metabolites to age-matched healthy controls (n=35) enrolled in a prospective cohort study in Lusaka, Zambia (NCT04010448 ClinicalTrials.gov, Lusaka, Zambia). Tryptophan levels were similar between groups, however RV-infected infants exhibited significantly lower levels of microbiota-derived indole metabolites, including IAA, IPA, tryptamine and tryptophol relative to healthy controls **(Fig. 5, Supplementary Fig. 5)**. While kynurenine itself showed no significant differences, its downstream metabolites, picolinic acid and xanthurenic acid were reduced in RV-infected infants **(Fig. 5, Supplementary Fig. 5).** These findings confirm that diminished microbiota-derived indole metabolites are associated with RV infection in a high-risk infant population, reinforcing the potential role of microbiota-derived indole metabolites in modulating RV infection and replication.

**Figure 5.**
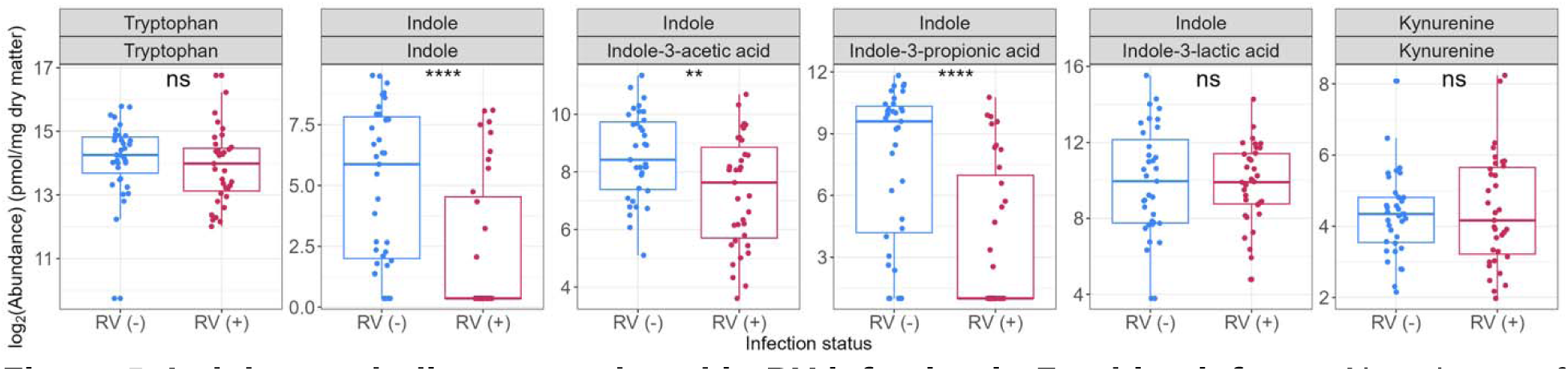
Indole metabolites are reduced in RV infection in Zambian infants. Abundance of select tryptophan metabolites in RV-infected infants (n=35) and age-matched non-infected counterparts (n=35). The bottom header of each panel shows the metabolite names, top header indicates the pathway classification (kynurenine or indole pathway). Boxplot box indicates first, second and third quartiles of data. P-values are based on Wilcoxon test with FDR correction (* p< 0.05; ** p< 0.01; *** p< 0.001, **** p< 0.0001).

## Discussion

In this study, we demonstrate that microbiota-derived indole metabolites associate with protection against rotavirus (RV) infection in both healthy adults and Zambian infants. Further, we show that indole-3-acetic acid (IAA) restricts RV replication through activation of the aryl hydrocarbon receptor (AhR) pathway using relevant human intestinal enteroids (HIE) models and a murine model of RV infection. Antibiotic-induced modulation of the gut microbiota in healthy adults led to reduced levels of fecal indole metabolites and associated with increased fecal RV shedding. Conversely, pre-treatment with IAA in HIEs and with the AhR agonist indole-3-carbinol (I3C) in mice directly suppressed RV replication. In line with these findings, Zambian infants with active RV infection exhibited lower indole metabolites levels compared to healthy age-matched controls. Together, these results suggest that microbiota-derived indole metabolites confer protection against RV infection through AhR-dependent mechanisms.

AhR is a ligand-activated transcription factor that responds to a wide range of environmental stimuli, regulating pathways critical in the maintenance of intestinal homeostasis, barrier function and mucosal immunity^50^. AhR is widely expressed across different cell types, including epithelial and endothelial cells as well as various immune cell populations^50^. In our study, IAA pre-treatment suppressed RV replication in HIEs devoid of immune cells, indicating that epithelial-intrinsic AhR activation is sufficient to confer antiviral effects. This was supported by RNA-seq analysis, which showed robust induction of AhR target genes, including Cyp1A1, confirming AhR pathway activation in these small intestine epithelial cells.

Several mechanisms may explain the AhR-mediated inhibition of RV replication. Our RNA-seq results indicated IAA pre-treatment led to upregulation of OAS1, an interferon-stimulated gene (ISG) downstream of the IFN pathway, suggesting that the IFN pathway may account for some of the observed reduction in RV replication following IAA treatment. Prior study have shown that exogenous IFN treatment limits RV replication in HIEs^42^. Our finding that AhR activation induces an IFN response contrasts with previous studies involving non-enteric viruses, where AhR activation suppresses type I IFN production and thereby promotes virus replication in various cell line models^51–53^. This discrepancy may be explained by differing model systems and viruses being studied, which potentially favor distinct IFN response. While previous studies primarily focused on type I IFN response, we suspect that type III IFN response is predominant in our model, as it is the main IFN response following RV infection in the intestinal epithelium^42,54,55^.

Alternative, possibly complementary, mechanisms by which IAA-induced AhR activation reduces RV replication in IECs are through the enhancement of gut barrier function. IAA has been reported to strengthen gut barrier function via both AhR-dependent and independent mechanisms^56–58^. Previous studies have demonstrated that improving gut barrier integrity can prevent recurrent RV infections and reduce disease severity in both gnotobiotic animal models^59^ and clinical studies in children^60^. Additionally, IAA has been shown to modulate mucin sulfation through an AhR-dependent pathway, contributing to a thickened mucus layer^57^. This mucin layer may serve as a physical barrier to RV^61^, reducing its ability to bind to IECs^17^, and thereby modulating host susceptibility to RV infection. Future study should evaluate barrier function and mucin availability in intestinal cells following IAA treatment and how that may affect RV attachment and host susceptibility to RV infection.

We also observed AhR mediated restriction of RV in our complex *in vivo* murine model of RV infection. In vivo, AhR activation may not only act on IECs, but also via immune cells. AhR agonism of innate lymphoid cells (ILCs) induces the production of interleukin-22 (IL-22)^62,63^, a cytokine critical for antiviral defense against RV^64,65^. Mice treated with AhR agonist I3C showed act in concert with type III IFN to promote antiviral ISGs expression, optimizing host defenses against RV infection^66^. Additionally, IL-22 enhances IECs turnover by promoting proliferation and extrusion of RV-infected cells, aiding in viral clearance^64,65^. Together, these mechanisms may explain the reduction in RV replication in mice treated with I3C. In line with our data, others have shown that antibiotic-treated mice, which exhibited reduced IL-22 expression, had increased RV replication^67^. Taken together, our findings and previous studies suggest that a disrupted or depleted gut microbiota may reduce levels of microbiota-derived AhR ligands, thereby impairing IL-22 production, and rendering the host more susceptible to RV infection.

Our findings contrasts with previous studies on non-enteric viruses, where AhR activation promoted the replication of zika virus in liver cells^52^ and coronavirus in lung epithelial cells^68^. A key distinction is that most prior studies used synthetic AhR agonists, host-derived kynurenine, or AhR knockout models^40^, whereas our study is the first to investigate the role of microbiota-derived AhR ligands in viral infection. We demonstrate that microbiota-derived AhR ligands, such as IAA, elicit distinct downstream effects compared to host-derived AhR ligands such as kynurenine. While RNA-seq data confirmed AhR activation with both kynurenine and IAA treatment, the downstream pathways they activated differed markedly. IAA induced genes in involved in antiviral defense such as ISGs, meanwhile kynurenine also showed strong upregulation of matrix metalloproteinases (MMP)1 expression (Figure 3C). MMPs are proteolytic enzymes involved in extracellular matrix remodeling and excess MMP activity can worsen disease pathology and pathogen infection by promoting inflammation and epithelial damage^69^, potentially explaining the lack of protection against RV infection with kynurenine treatment.

A key strength of our study is its translational approach, integrating evidence from adult and high-risk infant cohorts, with mechanistic validation in HIEs and murine model, demonstrating that indole metabolites consistently associate with protection against RV and providing the first evidence implicating the AhR pathway in enteric viral infection. The use of HIEs highlights that AhR activation in epithelial cells alone, in the absence of immune cells, is sufficient to suppress RV replication. Importantly, using these orthogonal approaches revealed notable differences between adult and infant HIEs following RV infection. Infant HIEs exhibited a weaker IFN response compared to adult HIEs, evidenced by upregulation of ISGs following RV infection in adult HIEs, but not infant HIEs. This aligns with previous findings that infant HIEs exhibit lower IFN response to the monovalent type 1 oral polio vaccine (mOPV1) and an overall lower innate epithelial immune response compared to adult HIE^70^. These findings highlight the importance of using multiple relevant models and accounting for age-specific immune responses when studying RV pathophysiology and developing novel RV treatments.

However, some limitations to our study remain. While we demonstrated that IAA activates AhR and suppress RV replication, the exact downstream mechanisms remain unclear and warrant further investigation. Additionally, our RNA-seq study focused on transcriptional changes at an early timepoint (5 hpi), possibly missing relevant mechanism arising later in infection. Second, the use of murine RV in our *in vivo* model may not fully recapitulate human RV infection, requiring further validation with human RV strains. Finally, our observational study in Zambian infants measured tryptophan metabolites at the time of infection rather than prior, and therefore confounding by infection itself cannot be ruled out.

Our findings suggest that depletion of microbiota-derived IAA may create a more permissive intestinal milieu for enteric virus infection, and that restoring AhR ligand levels through microbial, dietary or pharmacologic means could offer a protective strategies, particularly in vulnerable populations. In our study, oral I3C administration effectively reduced RV replication in mice, suggesting that dietary AhR agonists may be beneficial. Supporting this, prior work showed that a tryptophan-rich diet enhanced protection against RV infection and diarrheal disease in malnourished neonatal gnotobiotic piglets colonized with human infant fecal microbiota, via modulation of cytokine profiles and enhancement of RV-specific IgA antibody titers^71^. Future studies should focus on testing the efficacy of IAA and other (microbiota-derived) AhR agonists in protecting against RV infection in human populations, especially children living in high-risk settings.

In conclusion, by integrating observational data from an adult discovery cohort, *in vitro* and *in vivo* validation of primary findings as well as confirmation in an infant cohort, our study serves as a proof-of-concept that microbiota-derived metabolites can be leveraged for novel prevention and treatment strategies to reduce the burden of RV disease in vulnerable populations.

## Methods

### Adult challenge study

Fecal samples utilized in this study were obtained from a previous single-center, randomized (1:1:1), open-label, controlled study conducted at the Amsterdam University Medical Center, the Netherlands (METC 2015_045, NL 52510.018.15). The detailed study protocol has been described previously^7^. In short, 63 recruited adult volunteers were randomized into three antibiotic-treatment groups (n= 21 per group): control (no antibiotics), narrow-spectrum antibiotics (oral administration of vancomycin 500 mg three times a day), and broad-spectrum antibiotics (oral administration of vancomycin 500 mg and metronidazole 500 mg three times a day, ciproflaxin 500 mg twice a day). Participants received antibiotics treatment for 7 days, followed by a 48-hour wash-out period, before they were vaccinated with one dose of the oral rotavirus vaccine (ORV) Rotarix. ORV shedding was monitored through an ELISA assay to measure RV antigen level in fecal samples collected 1-7 days post vaccination. For this study, fecal samples collected 7 days post-antibiotics treatment but prior to ORV administration were used for metabolomics and metagenomics analysis.

### Zambian infant cohort

RV-positive samples (n=35) were a subset of fecal samples collected during a diarrhea surveillance study in Zambian children (ClinicalTrials.gov NCT04312906). Inclusion criteria included samples from infants aged 2 years or younger that tested positive for RV based on qPCR assay targeting the NSP3 gene using a Ct cut-off of <35. Age-matched RV negative samples (n=35) were selected from fecal samples collected during the follow-up period of the ROTA-biotic study (ClinicalTrials.gov NCT06882070) which was nested within a phase III randomized-controlled trial comparing the efficacy of a subunit P2-VP8 to ORV in Zambian infants (ClinicalTrials.gov NCT04010448). The ROTA-biotic study was approved in Zambia by the Zambian National Health Research Authority (NHRA) and the University of Zambia Biomedical Research Ethics Committee (UNZABREC) (Ref 1100-2020, August 21, 2020). The diarrhea surveillance study was approved by UNZABREC (ref 696-2020) and the NHRA. All included ROTA-biotic samples (controls) were non-diarrheal, tested negative for RV based on a qPCR assay, and had not received any antibiotic treatment prior to sample collection.

### Metabolomics analysis

Adult participants’ fecal samples: Global untargeted metabolomics analysis of the fecal samples was performed by the Metabolon, Inc (Durham, NC, USA) using a combination of four ultrahigh performance liquid chromatography tandem mass spectrometry (UPLC-MS/MS) methods to cover a wide spectrum of metabolites detection. All methods employed a Waters ACQUITY UPLC system coupled with a Thermo Scientific Q-Exactive high-resolution/accurate-mass spectrometer. A total of 1093 metabolites were identified and assigned to their respective super-pathway and sub-pathway according to Metabolon classification. Raw data were normalized by sample mass and re-scaled to set the median equal to 1. Missing values were imputed with the minimum observed value.

Zambian infants’ fecal samples: Targeted metabolomics to measure a curated list of tryptophan metabolites was performed according to a previously published protocol^36^. Tryptophan and kynurenine were quantified via HPLC using a colorimetric electrode assay (ESA Coultronics, MA, USA). Quantification was achieved using calibration curves generated with internal standards as a reference. Indole metabolites were quantified by UPLC-MS/MS using a Waters ACQUITY ultra performance liquid chromatography system that was coupled to a tandem quadrupole-time-of-flight (Q-TOF) mass spectrometer. Concentrations with a value below the lower limit of quantification (<LLOQ) were imputed as half of the minimum non-zero observed values.

To identify differential abundance metabolites between treatment groups, we performed Mann-Whitney U test with FDR correction for multiple testing using the R package omu^72^. Metabolites are considered differentially abundant when log2foldchange >1 or <-1, and p.adjusted <0.05.

### Cell line and virus strain

MA104 cells (African green monkey kidney epithelial cells), kindly provided by Dr. Erwin Duizer (National Institute for Public Health and the Environment, RIVM, the Netherlands), were used for virus culture. Cells were maintained in Eagle’s minimum essential medium (EMEM, Lonza) supplemented with 10% (v/v) heat-inactivated fetal bovine serum (FBS, Sigma-Aldrich, #F2442), 100 U/mL penicillin/streptomycin (Capricon scientific, #PSG-B), 1% (v/v) non-essential amino acids (Gibco, 11140-035) and 0.1% (v/V0 L-glutamine (Gibco, #25030-081). Cells were kept at 37°C, 5% CO_2_, 95% humidity and were passaged every 7 days using trypsin.

The human rotavirus WA strain (G1P8) was purchased from American Type Culture Collection (ATCC, Rotavirus A-VR-2018) and propagated on MA104 cells with no FBS supplementation. Rotavirus was harvested by freeze-thawing MA104 cells after they reached 90% cytopathic effect (CPE) following infection. The lysate from uninfected MA104 cells was used as a mock control.

### Human intestinal enteroids (HIE)

Adult HIE lines were commercially purchased from Lonza (751359, 751360) and Altis biosystem. Infant jejunal HIE lines (J1005, J1006, J1009) were generated at the Texas Medical Center Digestive Diseases Center Gastrointestinal Experimental Model Systems Core at Baylor College of Medicine. Tissue sample isolation protocol and characterization of infant lines have been published previously^70^. Both adult and infant lines were maintained as 3D HIE inside Matrigel (Corning, #356231) dome at 37°C, 5% CO_2_, and 95% humidity with IntestiCult™ Organoid Growth Medium Human (OGMh) (Stemcell™ Technologies, #06010) supplemented with 100 U/mL penicillin/streptomycin (Capricon scientific, #PSG-B). Medium was replenished every 2-3 days and HIEs were passaged every 3-5 days. For infection experiment, 3D HIEs were dissociated into single-cells and seeded onto Transwell® inserts (6.5 mm, 3.0 µm pore size, polycarbonate membrane, Corning #3415) coated with 0.02 mg/ml collagen type I (rat tail, Ibidi #50201) at a density of 1.0×10^5^ and 2.5×10^5^ cells per insert for adult lines and infant lines respectively. Cultures were differentiated in IntestiCult™ Organoid Differentiation Medium Human (ODMh) (Stemcell™ Technologies, # 100-0214) supplemented with rock inhibitor Y-27632 (Sigma, #Y0503) for the first three days. Afterwards, the medium was switched to ODMh without rock inhibitor and was replenished every other day until differentiation protocol was completed on day 14. Trans-epithelial electrical resistance (TEER) was measured after every medium change to monitor monolayer formation. Only inserts with TEER ≥ 300 Ω x cm^2^ were used for infection.

### Compounds preparation

Kynurenine (Sigma aldrich, # K8625), indole-acetic acid (Sigmal aldrich, #I3750) and AHR antagonist CH-223191 (Sigma aldrich, #C8124) were dissolved in DMSO and were stored at room temperature, −20°C, and 4°C respectively until further use.

### Infectivity assay

14-days differentiated HIE on Transwell inserts were treated with either 50 µM kynurenine, 500 µM IAA or 0.1% DMSO (vehicle control) in ODMh medium added to both apical and basal compartment for 24 hours before infections. Concentrations of compounds are based on physiological fecal levels in the Zambian infant cohort (with the assumption that fecal matter has a density of approximately 1g/mL). Rotavirus WA strain was activated by adding 10 μg/ml trypsin (Worthington biochemical, #9002-07-7) followed by incubation at 37°C for 1 hour. Virus was further diluted in ODMh medium supplemented with compounds and 0.25 mg/ml pancreatin to achieve an MOI of 1 FFU/cell. This inoculum was added to the basal compartment, and virus adsorption was allowed to occur for 4 hours at 37°C with 5% CO_2_. After adsorption, the Transwell inserts were transferred to a new 24-well plate containing fresh ODMh medium supplemented with compounds and 0.25 mg/ml pancreatin. Apical medium was replenished with ODMh supplemented with compounds, and the infection was continued overnight for another 20 hours.

### Viability assay

Cell viability was evaluated 48 hours post-incubation using ATP and LDH assay. Intracellular ATP level was measured using the 3D CellTiter-Glo Luminescent cell viability assay (Promega, #G9683) according to manufacturer’s protocol. LDH release was measured from apical supernatant collection using the LDH-Glo™ Cytotoxicity Assay (Promega, #J2381) according to manufacturer’s protocol. Luminescence was measured using an H1 synergy plate reader (BioTek).

### Animal experiment

All animal experiments were approved by the Washington University Institutional Animal Care and Use Committee. Wildtype C57BL/6 mice were originally purchased from Jackson Laboratory (Bar Harbor, ME) and bred and housed in Washington University in Saint Louis animal facilities under specific pathogen free conditions according to university guidelines. Mice were fed with standard rodent chow diet with ad libitum access to water. At 8-12 weeks of age, mice were randomized into two treatment groups, one group received daily oral gavage of I3C at the dose of 25 mg per kg body weight per day, while the other group received equal volume of PBS. 3 days after beginning I3C/PBS treatment, mice were orally inoculated with 10^4^ shedding dose 50 of murine rotavirus strain EDIM-Cambridge (ECRV). Treatment with either I3C or PBS was continued until day 7 post-ECRV infection, after which the mice were sacrificed, and tissue samples were collected and snapped frozen until further use. Daily fecal samples were collected to assess RV fecal shedding, which was quantified using RT-qPCR as previously described^73^.

### Fluorescence-focus assay (FFA)

Rotavirus titer was determined using the FFA method in MA104 cells. The monolayer from Transwell inserts were detached from the membrane by treatment with TrypLE (Gibco, #12605-010) for 10 min at 37°C. The cells were resuspended in serum-free EMEM and freeze-thawed to release intracellular virus. The resulting lysate was serially diluted with serum-free EMEM and added as an inoculum to confluent MA104 monolayers on 96-well plates. After a 1-hour adsorption period at 37°C, the inoculum was removed, and the cells were washed three times with serum-free EMEM. Infection was allowed to proceed overnight for 15 hours. The cells were then fixed with ice-cold methanol and stained with an anti-rotavirus polyclonal sheep primary antibody (1:300 dilution, Invitrogen, #PA1-85845) followed by an Alexa Fluor^TM^ 488-conjugated donkey anti-sheep secondary antibody (1:1000 dilution, Invitrogen, # A-11015). Rotavirus-positive cells were counted, and the results were reported as FFU/ml.

### RNA-sequencing

HIE monolayers were lysed in lysis buffer RLY from the Isolate II RNA Mini Kit (Bioline, #BIO-52073) and total RNA was extracted according to manufacturer’s protocol. Isolated RNA was sent to Macrogen, Inc (Netherlands) for sequencing. Any contaminating DNA was removed through DNAse treatment and RNA quality was checked using Agilent 2100 Bioanalyzer to ensure a RIN value of 8 or higher. Library preparation was performed using the Watchmaker mRNA library prep for poly-A selection, and sequencing was carried out on the NovaSeq X platform (Illumina) to produce 150 bp, paired-end reads. Raw sequences were quality-controlled with FastQC v0.11.7, trimmed with Trimmomatic 0.38. and mapped to the human genomes using HISAT2 version 2.1.0. Differential gene expression was determined by DEseq2 using Benjamini-Hochburg FDR corrected P-values and enrichment analysis was performed in Partek Flow (v.12.2.0). Consecutive data handling and plotting was performed using R (v. 4.4.0) R packages tidyverse (v.2.0.0) and ggplot2 (v.3.5.1). Volcano plots were generated with EnhancedVolcano (v. 1.22.0), heatmaps with ComplexHeatmap (v. 2.20.2).

### Statistical analysis

In the population studies, boxplots display the first, second and third quartiles of data, with whiskers representing the minimum and maximum values. Statistical significance was assessed using the Wilcoxon test with FDR correction for multiple testing. Correlation analysis between metabolite abundance and the area under the curve (AUC) of ORV shedding was performed using the Spearman Rank Correlation test. Statistical analyses were conducted using the R package ggpubr (version 0.6.0).

For the infectivity assay, each experimental condition was tested with six biological replicates (three adult and three infant HIE lines), with two technical replicates each. Statistical analysis was performed using Prism9 software (GraphPad). Statistical significance was assessed using the Kruskal-Wallis test with Dunn’s multiple comparisons test, where *p< 0.05, **p< 0.01, ***p< 0.001.

## Acknowledgement

We acknowledge and thank all staff, participants and guardians involved in the ROTA-biotic and diarrhea surveillance studies in Zambia. We also thank the participants from the adult challenge study in the Netherlands. All work with the adult HIE lines were supported and made possible by the OrganovirLabs at Amsterdam University Medical Centers. We are especially grateful to Dr. Carlemi Calitz, Dr. Adhitya Sridhar, Dr. Katja Wolthers and Prof. Dr. Dasja Pajkrt for their expertise and insightful discussion on the HIE study, and to Joep Korsten, Nina Johannesson and Gerrit Koen for their valuable technical assistance. The Texas Medical Center Digestive Diseases Center P30 DK056338 supported the generation of infant enteroids obtained from Baylor College of Medicine. We thank Antoine Lefevre and Patrick Emond at Université de Tours for performing the targeted metabolomic analysis. This research was supported by fundings from Amsterdam University Medical Centers and Amsterdam Institute for Infection and Immunity V.000332 (V.C.H and B.S); the Netherlands Organisation for Health Research and Development (ZonMw) VENI 09150161810022 (V.C.H.); National Institutes of Health (NIH) R01 AI173360 (S.R.,V.C.H and M.T.B).

## Authors contributions

N.I.W, B.S and V.C.H conceived and designed the study. N.I.W, Y.S, J.K, J.F, N.S and S.F performed the experiments. N.I.W, Y.S, J.K, N.P and G.J.P analyzed the data. C.C.L, M.C, M.S, C.C, S.S, K.C, B.T.N and D.H.K. managed and coordinated clinical study. M.S, S.R., M.T.B, B.S and V.C.H provided resources. M.S, S.R, D.M, M.T.B, B.S and V.C.H supervised the work. S.R.,M.T.B, B.S and V.C.H acquired funding. N.I.W and V.C.H wrote original draft. All authors reviewed and accepted the final contents of the manuscript.

