## Supplemental figures for "Microbiota-derived indole metabolites inhibit rotavirus infection *in vitro* and *in vivo* and in human infants"

**Supplementary figures**

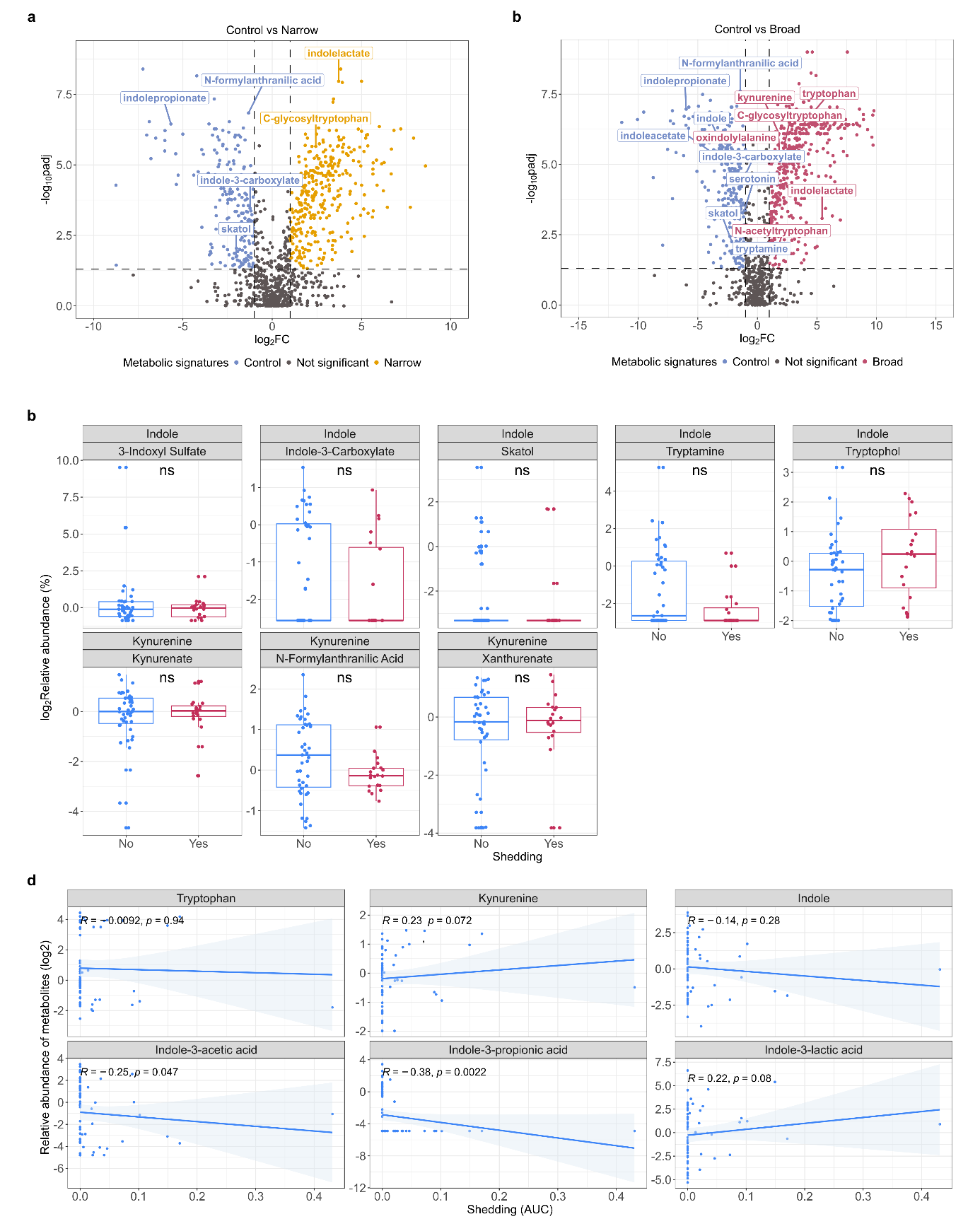

**Supplementary Figure 1. Associations between fecal tryptophan metabolites and fecal RV shedding in adult challenge study.** Volcano plot showing differentially abundant metabolites (log2foldchange >1 or <-1; p.adj <0.05, Mann-Whitney U test with FDR correction) in fecal samples, comparing **(a)** the control group with the narrow-spectrum group and **(b)** the control group and the broad-spectrum group. Metabolites belonging to the tryptophan group are labeled. **(c)** Relative abundance of other tryptophan metabolites in RV shedders (n=20) and non-shedders (n=43). The bottom header of each panel shows the metabolite names, top header indicates the pathway classification (kynurenine or indole pathway). Boxplot box indicates first, second and third quartiles of data. P-values are based on Wilcoxon test with FDR correction (ns: non-significant, * p< 0.05; ** p< 0.01; *** p< 0.001). **(d)** Spearman rank correlation analysis of the area under the curve (AUC) of RV shedding with the relative abundance of select tryptophan metabolites.

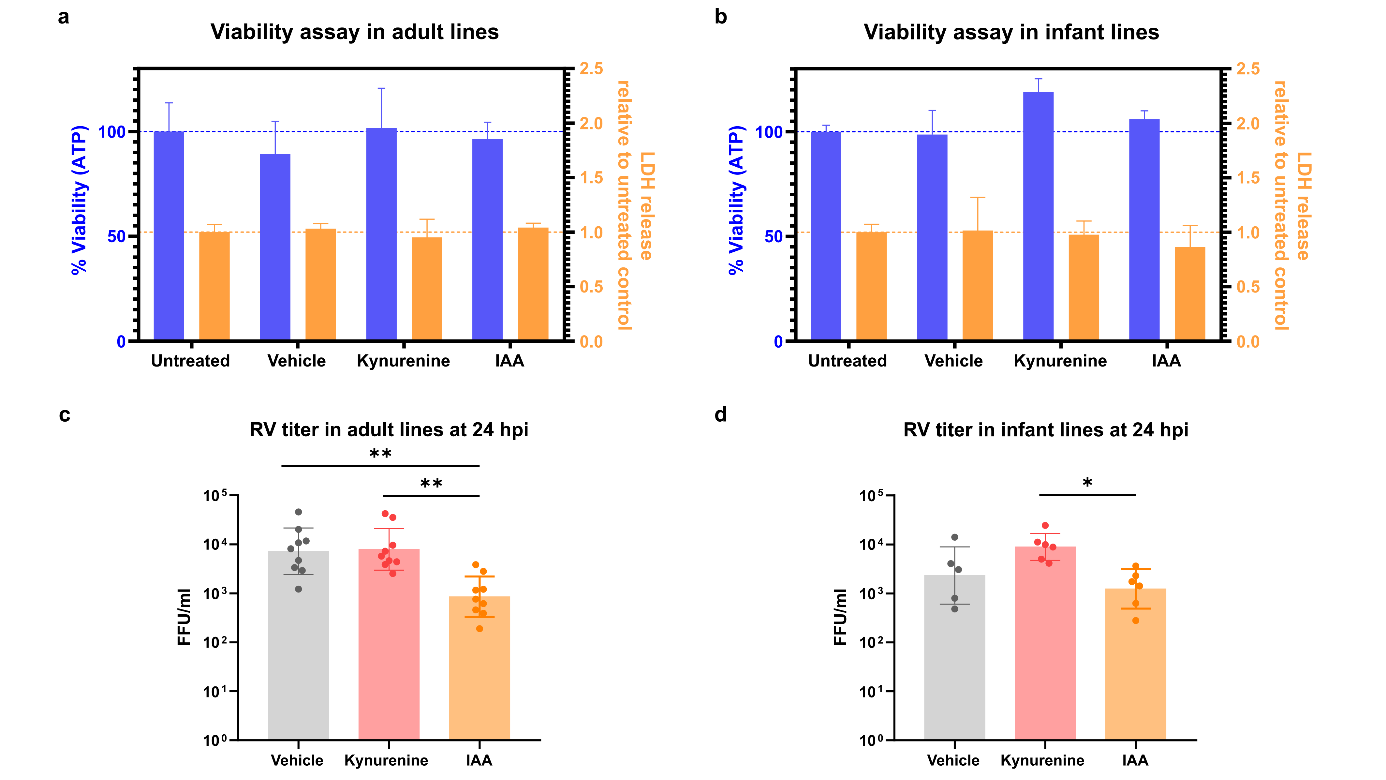

**Supplementary Figure 2. Viability and infectivity assay in adult and infant HIEs.** Percent cell viability in **(a)** adult and **(b)** infant HIE lines, assessed by ATP and LDH release. ATP levels are normalized to the untreated control, representing 100% viability (blue dashed line). LDH release is expressed as the relative change compared to the untreated control (orange dashed line). Production of RV infectious virus particles with 0.1% DMSO (vehicle), 50 µM kynurenine, or 500 µM IAA treatment, quantified at 24 hpi through fluorescence focus assay (FFA) in **(c)** adult and **(d)** infant HIEs. Data (geometric mean ± geometric SD) were collected from 3 adult and 3 pediatric HIE lines across 8 independent experiments. P-values are based on Kruskal-Wallis test with Dunn’s multiple comparisons test (* p< 0.05; ** p< 0.01; *** p< 0.001).

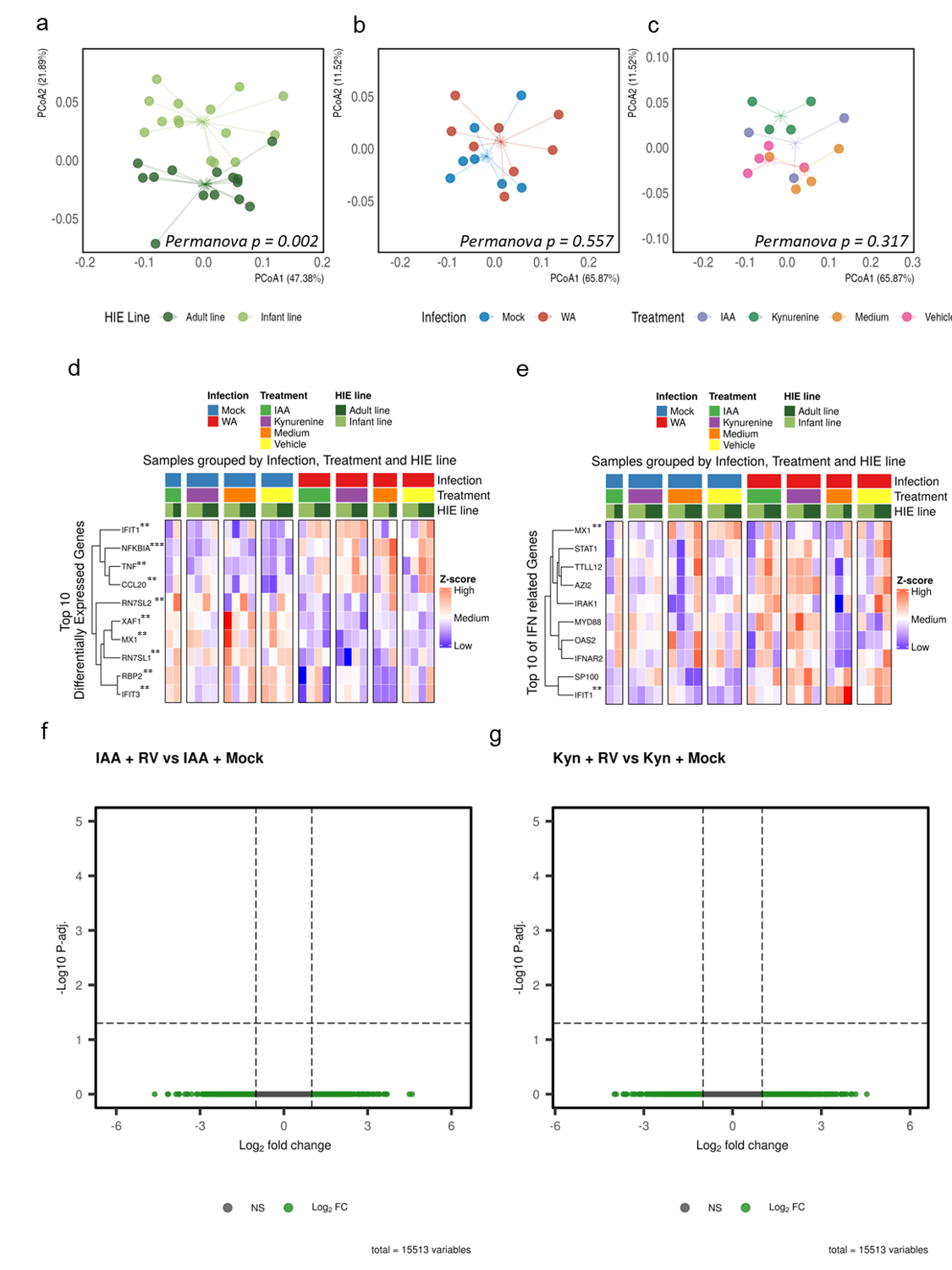

**Supplementary Figure 3. RNA sequencing analysis of adult and infant HIE treated with kynurenine and IAA. (a)** PC(o)A ordination indicating significant Bray-Curtis distance between adult and infant HIE lines. **(b)** PC(o)A ordination of infant HIE line indicating no significant Bray-Curtis distance of infection. **(c)** PC(o)A ordination of infant HIE line indicating no significant Bray-Curtis distance of treatment. **(d)** Heatmap of the top 10 most differentially expressed genes comparing mock- and RV-infected HIEs across treatment and HIE age group. **(e)** Similar to (d), but focusing only on IFN-related genes. P-values are based Wald t-test (* p< 0.05; ** p< 0.01; *** p< 0.001) Volcano plot showing differentially expressed genes (DEG) compared between **(f)** IAA-treated RV-infected and IAA-treated mock-infected in infant HIE line (n=2). **(g)** Kynurenine-treated RV-infected and Kynurenine-treated mock-infected in infant HIE line (n=2). DEGs were defined as log2foldchange >1 or <-1; p.adj <0.05. (Wald test with Benjamini-Hochberg FDR correction).

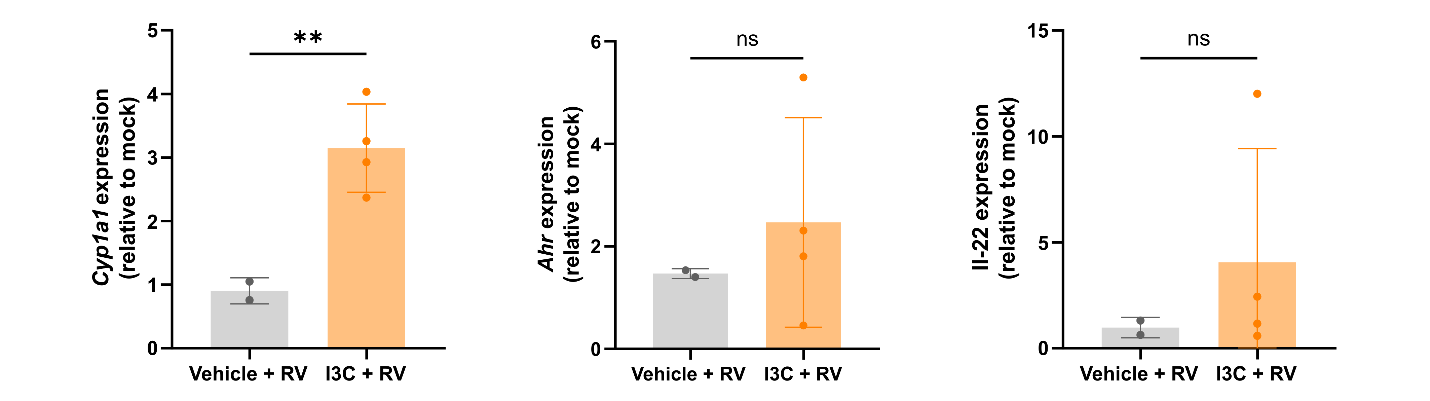

**Supplementary Figure 4. Gene expression analysis of intestinal tissue samples collected at 12 dpi.**

Expression of AhR target genes in intestinal tissue samples collected at 12 dpi. Gene expression levels were calculated relative to untreated uninfected mock control using the 2^-ddCt method. Data (mean ± SD are from 1 independent experiment with n=2-4 per group. P-values are based on Welch t-test (* p< 0.05; ** p< 0.01; *** p< 0.001).

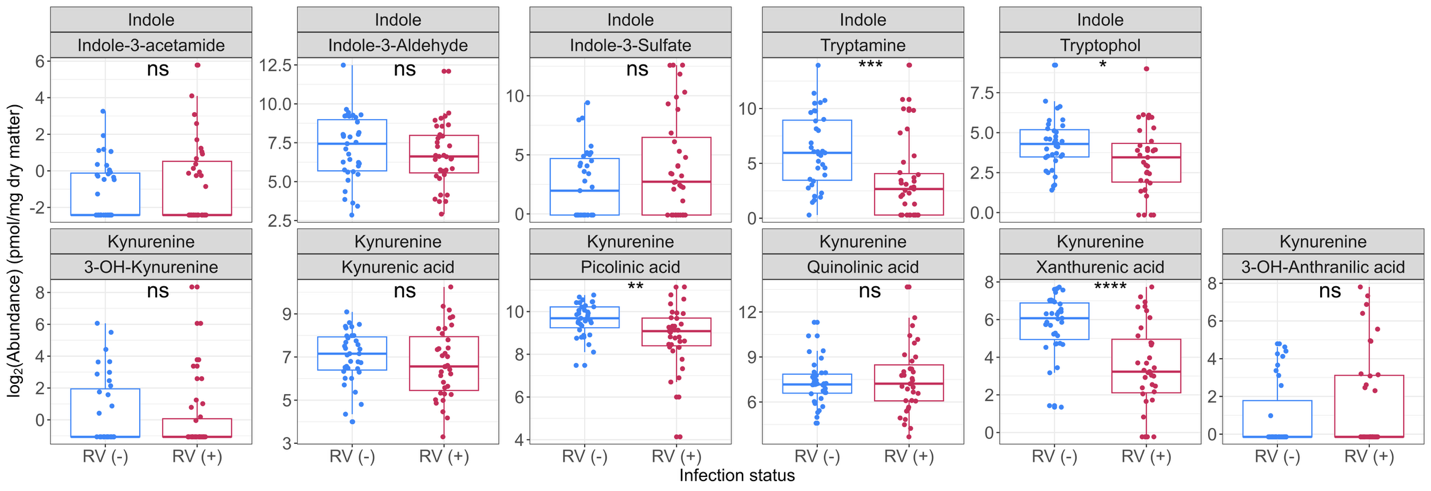
 **Supplementary Figure 5. Concentration of fecal tryptophan metabolites in RV-infected and non-infected Zambian infants.** Concentration of other tryptophan-derived metabolites in RV-negative (n=35) and RV-positive (n=35) infants. The bottom header of each panel shows the metabolite names, top header indicates the pathway classification (kynurenine or indole pathway). Boxplot box indicates first, second and third quartiles of data. P-values are based on Wilcoxon test with FDR correction (* p< 0.05; ** p< 0.01; *** p< 0.001, **** p< 0.0001).
